# Transcriptomic and epigenomic dynamics of honey bees in response to lethal viral infection

**DOI:** 10.1101/2020.08.14.251769

**Authors:** Hongmei Li-Byarlay, Humberto Boncristiani, Gary Howell, Jake Herman, Lindsay Clark, Micheline K. Strand, David Tarpy, Olav Rueppell

**Affiliations:** Department of Entomology & Plant Pathology, North Carolina State University, Raleigh, NC; Agricultural Research and Development Program, Central State University, Wilberforce, OH; Department of Biology, University of North Carolina at Greensboro, Greensboro, NC; Department of Entomology, University of Florida, Gainesville, FL; High Performance Cluster, Office of Information Technology, North Carolina State University, Raleigh, NC; High Performance Computing in Biology, University of Illinois at Urbana-Champaign, Urbana, IL; Army Research Office, Army Research Laboratory, Research Triangle Park, NC; W.M. Keck Center for Behavioral Biology, North Carolina State University, Raleigh, NC

**Keywords:** alternative splicing, transcriptome, DNA methylation, immune genes, pupa, IAPV, gene expression, comparative genomics

## Abstract

Honey bees (*Apis mellifera* L) suffer from many brood pathogens, including viruses. Despite considerable research, the molecular responses and dynamics of honey bee pupae to viral pathogens remain poorly understood. Israeli Acute Paralysis Virus (IAPV) is emerging as a model virus since its association with severe colony losses. Using worker pupae, we studied the transcriptomic and methylomic consequences of IAPV infection over three distinct time points after inoculation. Contrasts of gene expression and 5mC DNA methylation profiles between IAPV-infected and control individuals at these time points—corresponding to the pre-replicative (5 hr), replicative (20 hr), and terminal (48 hr) phase of infection—indicate that profound immune responses and distinct manipulation of host molecular processes accompany the lethal progression of this virus. We identify the temporal dynamics of the transcriptomic response to with more genes differentially expressed in the replicative and terminal phases than in the pre-replicative phase. However, the number of differentially methylated regions decreased dramatically from the pre-replicative to the replicative and terminal phase. Several cellular pathways experienced hyper- and hypo-methylation in the pre-replicative phase and later dramatically increased in gene expression at the terminal phase, including the MAPK, Jak-STAT, Hippo, mTOR, TGF-beta signaling pathways, ubiquitin mediated proteolysis, and spliceosome. These affected biological functions suggest that adaptive host responses to combat the virus are mixed with viral manipulations of the host to increase its own reproduction, all of which are involved in anti-viral immune response, cell growth, and proliferation. Comparative genomic analyses with other studies of viral infections of honey bees and fruit flies indicated that similar immune pathways are shared. Our results further suggest that dynamic DNA methylation responds to viral infections quickly, regulating subsequent gene activities. Our study provides new insights of molecular mechanisms involved in epigenetic that can serve as foundation for the long-term goal to develop anti-viral strategies for honey bees, the most important commercial pollinator.

**Author Summary:** Honey bees, the most important managed pollinators, are experiencing unsustainable mortality. Israeli Acute Paralysis Virus (IAPV) causes economically important disease in honey bees, and it is emerging as a model system to study viral pathogen-host interactions in pollinators. The pupation stage is important for bee development but individuals are particularly vulnerable for parasitic mite infestations and viral infections. Currently, it is unclear how honey bee pupae respond to this virus. However, these responses, including gene expression and DNA methylomic changes, are critical to understand so that anti-viral genes can be identified and new anti-viral strategies be developed. Here, we use next-generation sequencing tools to reveal the dynamic changes of gene expression and DNA methylation as pupae succumb to IAPV infections after 5, 20, and 48 hours. We found that IAPV causes changes in regions of DNA methylation more at the beginning of infection than later. The activity of several common insect immune pathways are affected by the IAPV infections, as are some other fundamental biological processes. Expression of critical enzymes in DNA methylation are also induced by IAPV in a temporal manner. By comparing our results to other virus studies of honey bees and fruit flies, we identified common anti-viral immune responses. Thus, our study provides new insight on the genome responses of honey bees over the course of a fatal virus infection with theoretical and practical implications.

## Introduction

Infectious disease results from the breakdown of an organism’s normal physiological state because of the presence or proliferation of a pathogen. This disruption can be molecularly characterized by transcriptome profiling to permit a systemic understanding of host-pathogen interactions, particularly in combination with other system-level approaches [1-3]. Transcriptomic and epigenomic changes in response to pathogen infections are important to understand because they are interdependent but relate to different temporal dynamics.

DNA methylation of Cytosine (5mC) is a central epigenetic regulator of gene expression and alternative splicing [4, 5], affecting diverse aspects of organismal function and disease [6]. Epigenetic programming may also be the target of host manipulations by pathogens [7] and affect host defenses and susceptibility to infection [8, 9]. The virus induced changes in host immune response and gene expression via DNA methylation is still a new study field. It is understudied whether changing host DNA methylation such as gene expression of certain antiviral immune genes is a main mechanism utilized by DNA or RNA viruses to manipulate immune responses. Genome-wide analyses of methylome and transcriptome indicated that the infection of DNA tumor virus induced hypermethylation of host genes during viral persistence [9]. In plants, DNA hypomethylation was reported after 24 hours of viral infection [10].

Honey bees (*Apis mellifera* L) were the first insects in which a fully functional 5mC DNA methylation system was discovered [11, 12]. Social hymenopterans, such as bees, wasps, and ants, have low levels of DNA methylation [13-15]. The methylated 5mC sites can be one of three different kinds (CpG dinucleotides, CHG, and CHH, H represents A, T or C) [16]. The CpG is the most common kind in honey bees. CpG DNA methylation in insects occurs predominantly in gene bodies, associated with histone modifications and alternative splicing [4, 5, 16-20]. Its phenotypic consequences range from behavioral plasticity [21, 22] to alternative development trajectories [12, 23] and disease responses [24]. Especially the study from Herb et al [21] showed gene expression differences related to differentially methylated regions (DMRs) and also suggests wider effect because many DMRs are related to transcription factors, in which the alternative splicing may affect their function. Despite this breath of effects, few studies have investigated the role of CpG DNA methylation in insect development, immune system, and disease [25-28]

Honey bees are the most important managed pollinator in agriculture and crop production [29]. However, recent declines in honey bee health have led to unsustainably high annual colony losses in the apicultural industry [29, 30] for which multiple, potentially interacting factors are likely the cause [31, 32]. One major contributor to widespread colony mortality is pathogenic viral infection. Many of the most important viruses are either associated with or can be directly vectored by parasitic *Varroa* mites [33-36]. Pathogenic viral stressors affect the morphology, physiology, behavior, and growth of honey bees at different developmental stages [31, 33, 37, 38], ultimately leading to reduced colony productivity and mortality. The developmental pupal stage is critical for many viral diseases because the *Varroa* mite feeds on the pupae for a prolonged time [35].

Honey bees employ a combination of individual- and group-level defenses against pathogens [39]. The main innate immune pathways of insects are present in honey bees and respond to virus infections [40, 41], including the Toll, Janus kinase and Signal Transducer and Activator of Transcription (JAK-STAT), Immune Deficiency (Imd), c-Jun NH2-Terminal Kinase (JNK) pathways, antimicrobial peptides (AMPs), endocytosis, and phagocytosis mechanisms [40]. Activation of some immune pathways, such as the RNAi mechanism, can lead to increased virus resistance [42]. Thus, we understand that some honey bee genes respond to virus infection but a systematic characterization of the temporal dynamics of the immune responses and the general transcriptome and methylome have not been investigated, although such studies can yield important information [43].

Israeli Acute Paralysis Virus (IAPV) has been associated with colony losses [44-46], particularly during the winter [47]. IAPV belongs to the picornavirus-like family Dicistroviridae, that includes single-strand, positive sense RNA viruses that are pathogenic to a range of insects [48]. IAPV is relatively common in honey bees and affects all life history stages and castes [47]. When cells are infected, the positive strand of RNA is released into the cytosol and translated by the host ribosomes. Then the assembled complexes for viral replication synthesize negative-sense RNA from which more descendant positive strands of the RNA are produced. These are assembled with viral proteins into virions that are then released to infect other cells [47, 49]. Covert infections persist through vertical and horizontal transmission, but IAPV can also readily cause paralysis and death of infected individuals [50]. Acute infections of IAPV profoundly affect cellular transcription at the pupal stage [51], and microarray analyses of larvae and adults indicate significant and stage-specific responses to IAPV infection [47]. Parallel transcriptomic and epigenomic profiling of adult IAPV infection identified some additional molecular responses, including expression changes in immune genes and epigenetic pathways [24].However, little overlap of identified responses within and among studies leave the molecular characterization of IAPV infection in honey bees unresolved, and more detailed studies of dynamic responses in the time-course of IAPV infection are needed.

Here, we experimentally test the hypothesis that viral infections can affect not only transcriptomic profiles but also induce DNA methylomic changes in honey bee pupae. Specifically, we compared the complete transcriptome and methylome profiles across three distinct phases of a lethal IAPV infection. Furthermore, we tested if our results indicate significant overlap of immune genes with fruit flies as well as other stages of honey bees under IAPV viral infections.

## Results

### Temporal viral infections cause transcriptomic changes in gene expressions

The transcriptomes of whole honey bee pupae injected with PBS or infected with IAPV were compared at three time points: 5h, 20h, and 48h post-injection (Figure 1). On average, IAPV-infected pupae yielded slightly more RNA-seq reads than the corresponding PBS controls (5h: IAPV = 46,001,777 vs. PBS = 41,666,177; 20h: IAPV = 47,915,086 vs. PBS = 42,429,737; and 48h: IAPV = 42,649,987 vs. PBS = 39,248,297). Coverage thus varied among individual samples from 41x to 65x (Supplemental Table 1). Differentially expressed genes (DEGs) increased with disease progression over time from 432 after 5h to 5,913 after 20h and 5,984 after 48h (Supplemental Table 2).

**Fig. 1:**
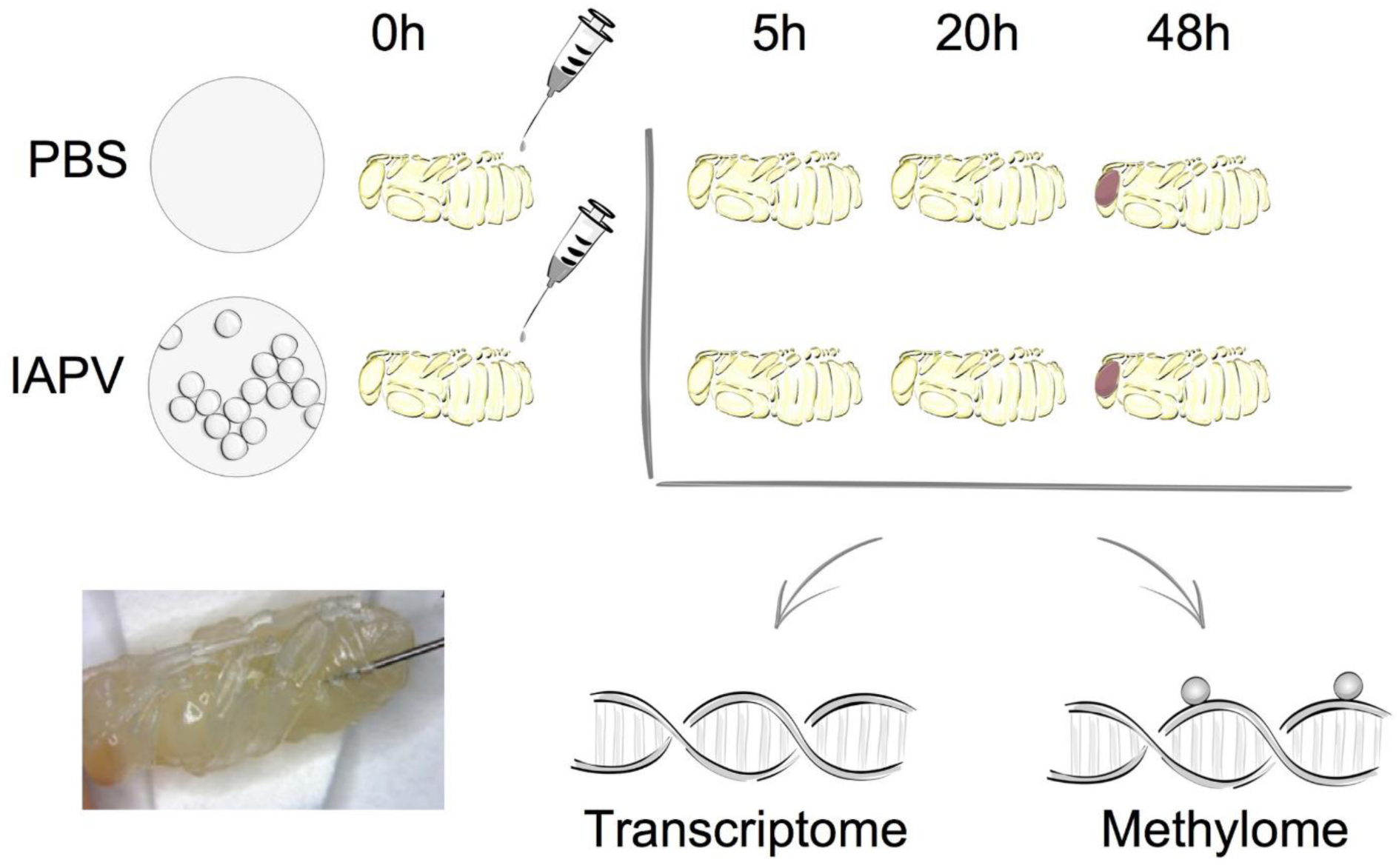
Overall experimental design. The two experimental groups included either sham treatment (PBS injection) or Israeli Acute Paralysis Virus (IAPV) inoculations via injection. At each investigated time point (5h, 20h, or 48 h post-injection) three biological replicates were collected, each consisting of one whole pupa that was individually processed for further transcriptomic and methylomic analyses.

Relative to the PBS control, IAPV-infected pupae had 198 genes significantly up-regulated and 234 genes significantly down-regulated after 5h; 2,738 genes were significantly up-regulated and 3,175 genes significantly down-regulated after 20h; and 3,021 genes significantly up-regulated and 2,963 genes significantly down-regulated after 48h (Supplemental Figures 1-3). DEG overlap between all significantly up- or down-regulated genes was significant in all pairwise directional comparisons (up- and down-regulated genes separately) among the three time points (Figure 2; up-regulated 5h vs 20h: p < 0.05; all other tests: p < 0.001). For the >8-fold DEGs, significant overlap was found for all pairwise comparisons among the IAPV up-regulated gene sets (p < 0.001) and also for the down-regulated gene sets at 20h and 48h (p < 0.001) (Supplemental Table 2).

**Fig. 2:**
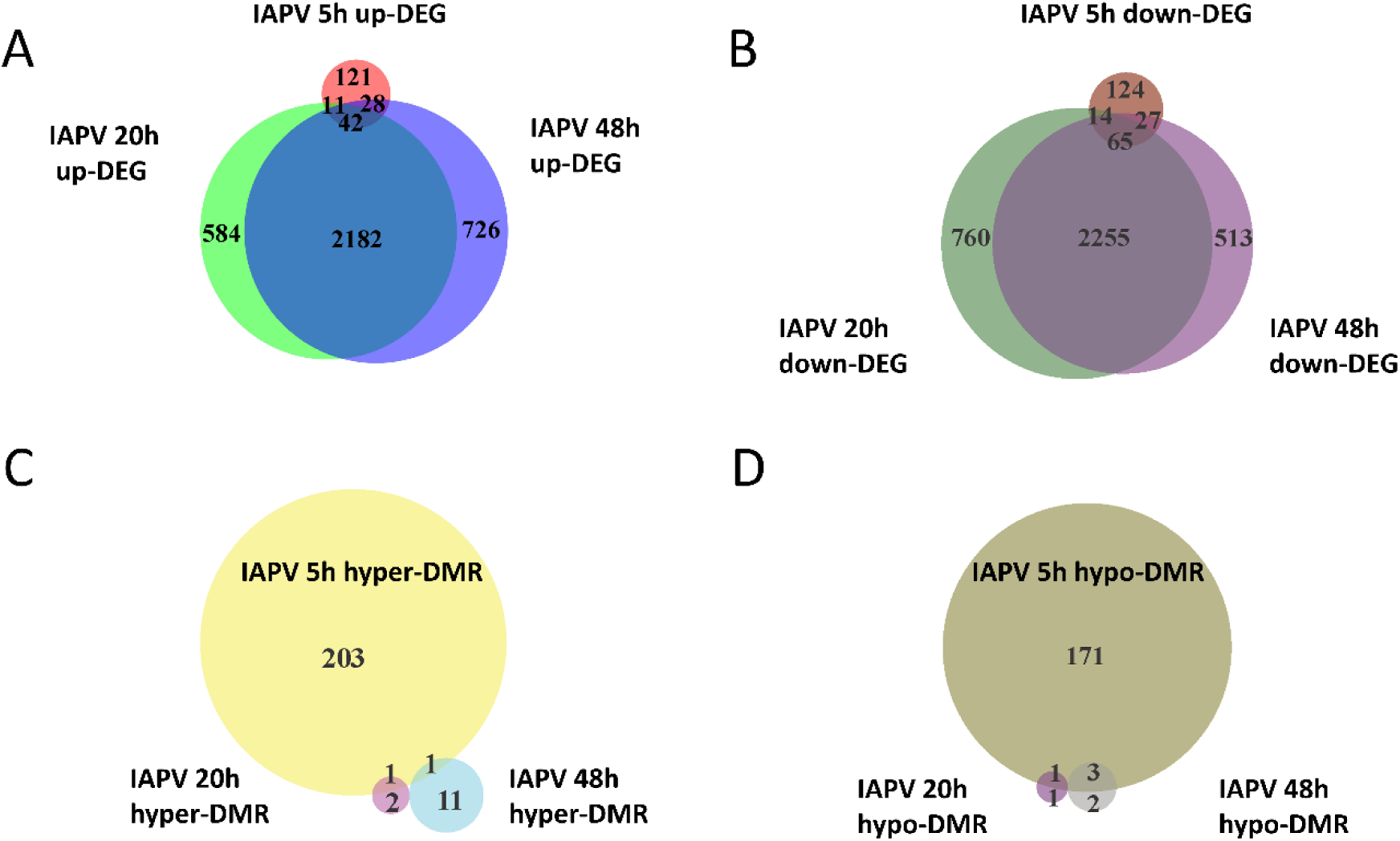
Directional overlap of differentially regulated genes among time points. (A) Venn diagram showing genes that were differentially expressed as up-regulated (FDR < 0.05). Number of DEGs only in post IAPV 5h, 20h, and 48h are 121, 584, and 726. Number of genes between IAPV 5h and 20h only overlap is 11. Number of DEGs between IAPV 5h and 48h only overlap is 28. Number of DEGs between IAPV 20h and 48h only overlap is 2182. Number of DEGs from IAPV 5h-20h-48h overlap is 42. (B) Venn diagram showing genes that are differentially expressed as down-regulated (FDR < 0.05). Number of DEGs only in post IAPV 5h, 20h, and 48h are 124, 760, and 513. Number of DEGs between IAPV 5h and 20h only overlap is 14. Number of DEGs between IAPV 5h and 48h only overlap is 27. Number of DEGs between IAPV 20h and 48h only overlap is 2255. Number of DEGs from IAPV 5h-20h-48h overlap is 65. (C) Venn diagram showing genes that are differentially methylated (percent methylation difference larger than 10%, q < 0.01) as hyper-methylation among three time points (5h, 20h, and 48h after IAPV infection). Number of DMRs only in post IAPV 5h, 20h, and 48h are 203, 2, and 11. Number of DMRs between IAPV 5h and 20h only overlap is 1. Number of DMRs between IAPV 5h and 48h only overlap is 1. (D) Venn diagram showing genes that are differentially methylated (percent methylation difference larger than 10%, q < 0.01) as hypo-methylation among three time points (5h, 20h, and 48h after IAPV infection). Number of DMRs only in post IAPV 5h, 20h, and 48h are 171, 1, and 2. Number of DMRs between IAPV 5h and 20h only overlap is 1. Number of DMRs between IAPV 5h and 48h only overlap is 3.

### Gene enrichment and functional analyses

To identify IAPV-induced expression patterns across all genes regardless of significance threshold, GO Mann-Whitney U tests and a Weighted Gene Co-expression Network Analysis were performed on the expression differences between IAPV infected and control samples at three time points. Post 5h of IAPV infection, multiple biological process GO terms were observed with upregulation of RNA processing (ncRNA, tRNA, and rRNA) as the most significant GO terms (p < 0.001). Only one term was significantly downregulated at this time point: “small GTPase mediated signal transduction”. 20h post IAPV infection, no GO terms passed the strictest significance threshold (p < 0.001) for upregulation, and GO term “translation” was the most significantly downregulated (p < 0.001). Finally, in post 48h IAPV infected samples, there was still no term passing the strictest significance in upregulation, and GO terms of “translation”, “homophilic cell adhesion”, and “aminoglycan metabolic process” were the most significantly downregulated (Figure 3). As to molecular function, significant GO terms associated with downregulation in post 20h and 48h IAPV infected samples were “structural molecules”, “structural constituent of ribosome”, “structural constituent of cuticle”, and “chitin binding” (Supplemental Figure 4). As to cellular component, significant GO terms associated with downregulation in post 20h and 48h IAPV infected samples were “ribosomal subunit”, “ribonucleoprotein complex”, and “intracellular non-membrane organelle” (Supplemental Figure 4). Although numerous other terms were less significantly enriched in up- or down-regulated genes, no other GO terms surpassed the most stringent significance criterion.

**Fig. 3:**
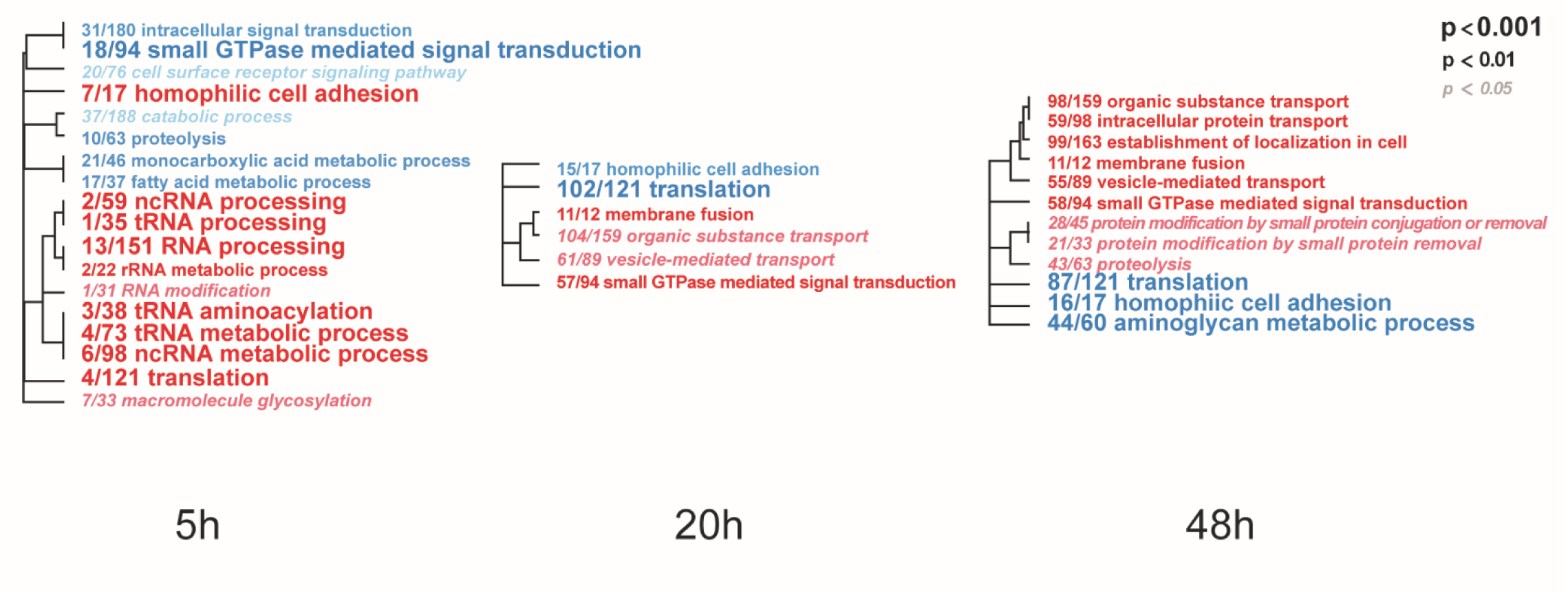
Overall trends in biological processes affected by progressing IAPV infection. Gene Ontology Mann-Whitney U tests of the expression differences between IAPV and control samples at the three time points included all data points regardless of their individual level of statistical significance. Red and blue fonts indicate GO terms that are enriched in up- and down-regulated DEGs in IAPV relative to control samples, respectively. Font size indicates the level of significance for the particular GO term and clustering is based on empirical GO-term similarity in our dataset.

### Viral and ribosomal RNA content

Four transcripts (IAPV, Deformed Wing Virus (DWV), Large Sub-Unit rRNA (LSU rRNA), & Small Sub-Unit rRNA (SSU-rRNA)) accounted for 63.5% of all sequenced reads (Supplemental Figure 5). As expected, a large number of IAPV reads was observed in the IAPV-infected pupae, with the proportion being similarly high at 20h (86% ave) and 48h (90% ave). Additionally, five of nine control samples (all time points included) had a large proportion of reads mapping to DWV.

### Comparative transcriptomic analyses of immune genes with other pathogen infection in honey bees and Drosophila

A comparison with a previously published list of 186 immune genes from Evans et al (2005) [41] showed that 1 and 89 of them were represented in our 5h, 20h/48h DEG lists respectively (Figure 4A, Supplemental Tables 3 and 4). The overlap between immune genes and our DEGs was significant for up-regulated DEGs at 5h and down-regulated DEGs at 20h (Supplemental Table 3 and 4). Comparison of our data with IAPV-induced gene expression changes in adult honey bees in Galbraith et al [24] revealed 6 and 358 genes overlapped, indicating a significant directional overlap of our up- and down-regulated DEGs at 5h and 20h/48 h with DEGs in adults (Figure 4C, for specific gene names, see Supplemental Table 3 and 4). When compared to immune genes associated with anti-viral responses in *Drosophila* [52], 11, 212, and 219 genes overlapped in our 5h, 20h, and 48h DEG lists (Supplemental Table 5), representing significant directional overlap of our up- and down-regulated DEGs at 20h and 48h (for specific gene names see Supplemental Table 5)

**Fig. 4.**
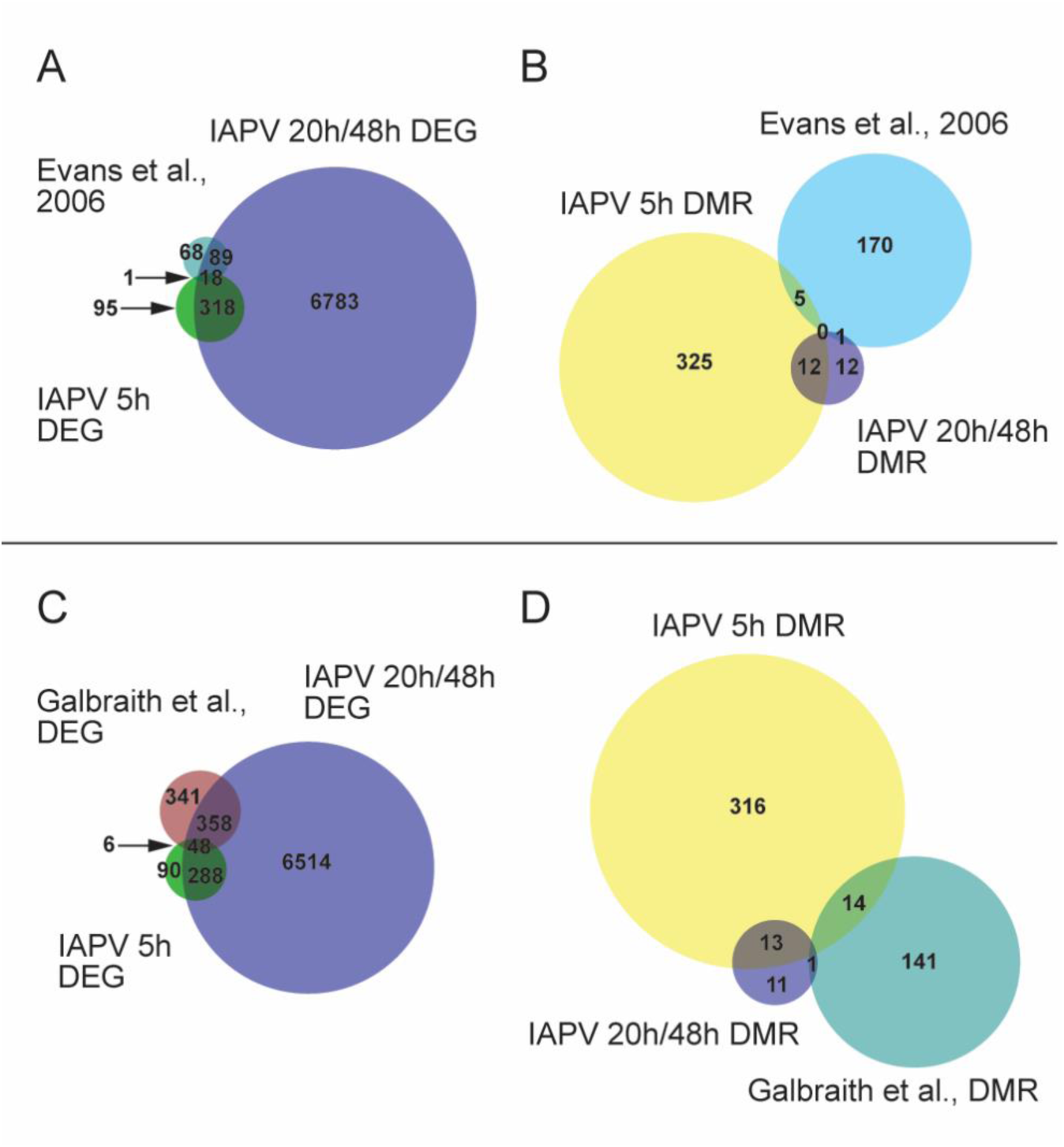
Comparisons or our results to other studies revealed significant overlap. (A) Venn diagram showing overlap of genes that are differentially expressed (FDR < 0.05; p < 0.05). Number of DEGs (differentially expressed genes) only in IAPV 5h, 20h/48h, or Evans et al 2006 immune gene list is 95, 6783, or 68. Number of DEGs overlap between IAPV 20h/48h and Evans et al is 89. Number of DEGs overlap between IAPV 5h and 20h/48h is 318. Number of DEGs overlap between IAPV 5h and Evans et al is 1. (B) differentially methylated (> 10%, p < 0.05) among three time points (5h, 20h, and 48h after viral infection) and immune gene list from Evans et al [34]. Number of DMRs (differentially methylated regions) only in IAPV 5h, 20h/48h, or Evans et al 2006 immune gene list is 325, 12, or 170. Number of DMRs overlap between IAPV 20h/48h and Evans et al is 1. Number of DMRs overlap between IAPV 5h and 20h/48h is 12. Number of DMRs overlap between IAPV 5h and Evans et al is 5. (C) Venn diagram showing overlap of DEGs (FDR< 0.05; p < 0.05) compared to Galbraith et al. Number of DEGs only in IAPV 5h, 20h/48h, or Galbraith et al is 90, 6514, or 341. Number of DEGs overlap between IAPV 20h/48h and Galbraith et al is 358. Number of DEGs overlap between IAPV 5h and 20h/48h is 288. Number of DEGs overlap between IAPV 5h and Galbraith et al is 6. (D) differentially methylated (> 10%, p < 0.05) among three time points (5h, 20h, and 48h after IAPV infection) and immune gene list from Galbraith et al DMR list. Number of DMRs (differentially methylated regions) only in IAPV 5h, 20h/48h, or Galbraith et al DMR list is 316, 11, or 141. Number of DMRs overlap between IAPV 20h/48h and Galbraith et al is 1. Number of DMRs overlap between IAPV 5h and 20h/48h is 14. Number of DMRs overlap between IAPV 5h and Galbraith et al is 13.

### Temporal dynamics of genes and molecular pathways based on transcriptomic data

Analyses of the three infection time points revealed dramatic changes of gene expression levels from several key metabolic and immune pathways, including lipid metabolism, the Jak-STAT pathway, and AMPs. With regards to lipid metabolism, the expression of the gene *Cyp6as5* was significantly increased in the infected samples from 5h to 20h, and from 20h to 48h, similarly to 9 more genes that displayed similar expression profiles (Supplemental Figure 7).

In the Jak-STAT immune pathway, the expression of gene *SoS* was significantly increased in the infected samples from 20h to 48h (Supplemental Figure 6). Nine more genes displaying similar expression profiles as *SoS* after viral infection were identified (Supplemental Figure 7). Among the AMP genes, the expression of gene *defensin-1* was significantly increased in the infected samples from 5h to 20h, and from 20h to 48h (Supplemental Figure 6). The gene expression of EGFR and a group of other genes were transiently downregulated from 5h to 20h post-infection and from 20h to 48h post-infection by IAPV (Supplemental Figure 7). Gene *GB55029* (uncharacterized) showed an increase from 5h to 20h infection, but then a significant decrease of expression from 20h to 48h infection (Supplemental Figure 7).

### Differentially spliced genes

Number of genes with significant isoform switches in samples 5h, 20h, and 48h post IAPV infection were 193, 620, and 990 (Supplemental Figure 8). Number of significant differential transcripts usage (DTU) or alternative splicing is much higher post 20h and 48h of IAPV infection (Supplemental Figure 9). All the splicing events are categorized in Table 1. Among all the categories, there are significant trends of DTU to be in shorter 3’ UTRs, longer 5’ UTRs, Exon gain, Transcription Start Site (TSS) more downstream in 20h and 48h IAPV post infection treatments (Figure 5).

**Table 1.** Isoform switch consequences investigated. Numbers of significant genes are shown.

| Category | Consequence | Description | Post 5h | Post 20h | Post 48h |
| --- | --- | --- | --- | --- | --- |
| 1 | Tss | Change in transcription start site | 120 | 407 | 602 |
| 2 | Tts | Change in transcription termination site | 21 | 169 | 246 |
| 3 | Last exon | Last exon changed | 9 | 97 | 130 |
| 4 | Intron retention | Difference in intron retention | 26 | 100 | 186 |
| 5 | Intron structure | Different exon-exon junctions used | 160 | 552 | 875 |
| 6 | Exon number | Different number of exons | 86 | 373 | 627 |
| 7 | ORF seq similarity | Jaccard similarity of AA sequences below 0.9 | 47 | 211 | 281 |
| 8 | ORF genomic | Change in genomic position of ORF | 97 | 382 | 540 |
| 9 | 5 utr length | Difference in length of 5' UTR | 131 | 433 | 653 |
| 10 | 3 utr length | Difference in length of 3' UTR | 27 | 190 | 284 |
| 11 | NMD status | Change in sensitivity to nonsense-mediated decay | 5 | 45 | 76 |
| 12 | Coding potential | Change in coding potential probability, above or below 0.7 | 5 | 30 | 36 |
| 13 | Domains identified | Change in which protein domains were identified | 19 | 96 | 119 |
| 14 | Domain length | Change in length of overlapping domains | 3 | 20 | 26 |
| 15 | IDR identified | Difference in presence of intrinsically disordered regions | 19 | 112 | 159 |
| 16 | IDR length | Difference in length of intrinsically disordered regions | 5 | 38 | 45 |
| 17 | IDR type | Difference in presence of binding site in IDR | 7 | 42 | 63 |
| 18 | Signal peptide identified | Change in presence of signal peptide | 1 | 8 | 14 |

**Fig. 5.**
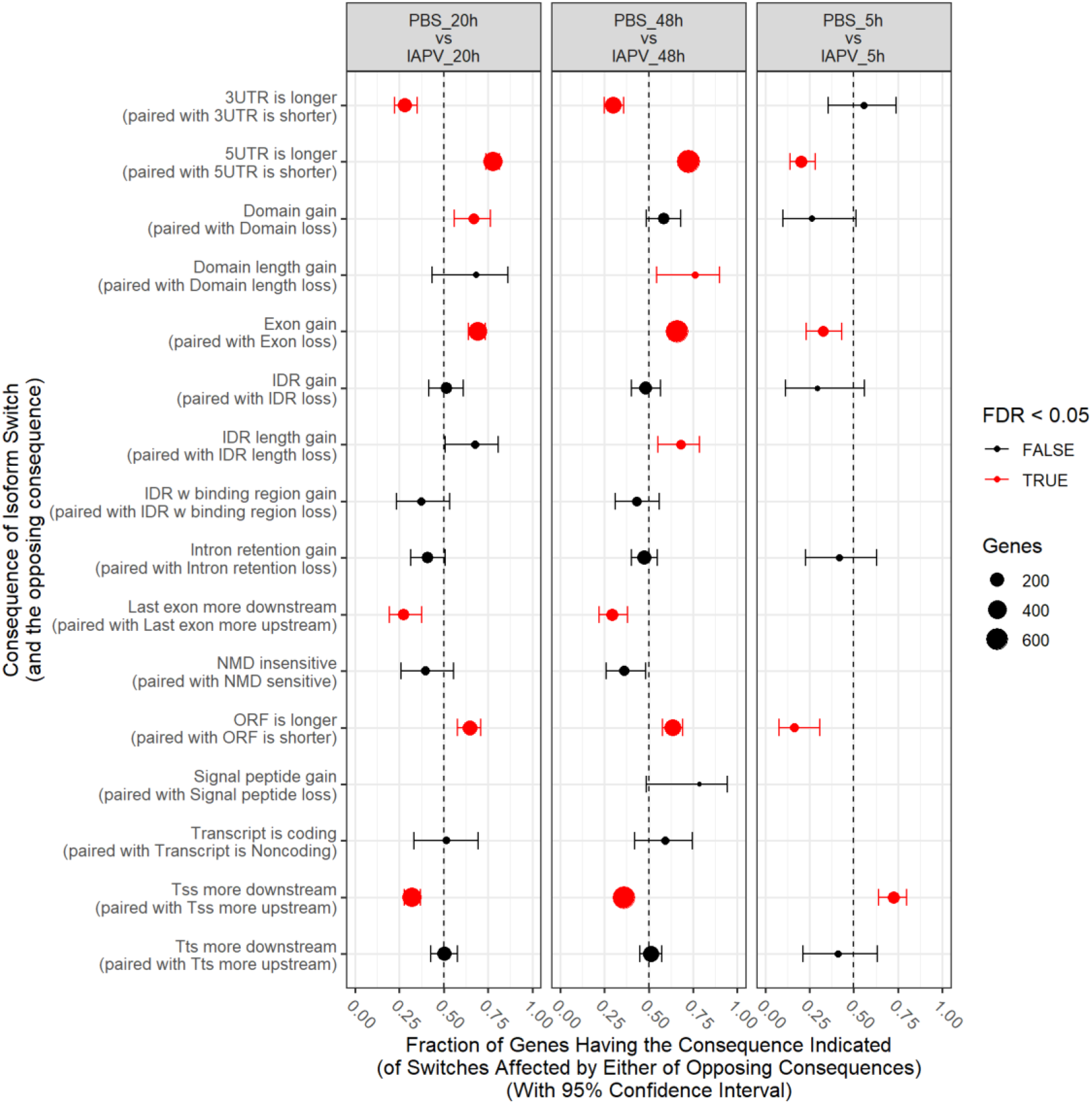
Significant trends in consequences of isoform switches or alternative splicing were observed, and were generally similar at 20h and 48h, often with opposite consequences at 5h. Fraction of genes having the consequence indicated with 95% Confidence Interval. UTR-untranslated region, ITR-Inverted Terminal Repeat, ORF-open reading frame, NMD-nonsense-mediated decay, TSS-transcription start site, TTS-transcription terminal site.

### Enzymes involved in DNA methylation

Gene expression differences were also identified in the DNA methylation system, specifically the two important enzymes, *DNA methyl-transferase 1* (*DNMT1*) and *3* (*DNMT3*). When compared between IAPV and control samples, *DNMT1* (*GB47348*) was significantly down-regulated 1.5 fold (p<0.001) at 20h post-infection, and 1.7 fold (p<0.001) at 48h post-infection (Figure 6). Similarly, *DNMT3* (*GB55485*) was significantly down-regulated 1.7 fold (p<0.001) in 20h post-infection samples, and 2.7 fold (p<0.001) in 48h post-infection samples (Figure 6). There was no significant difference of gene expression between IAPV and controls post 5h post-viral infection (Figure 6).

**Fig. 6:**
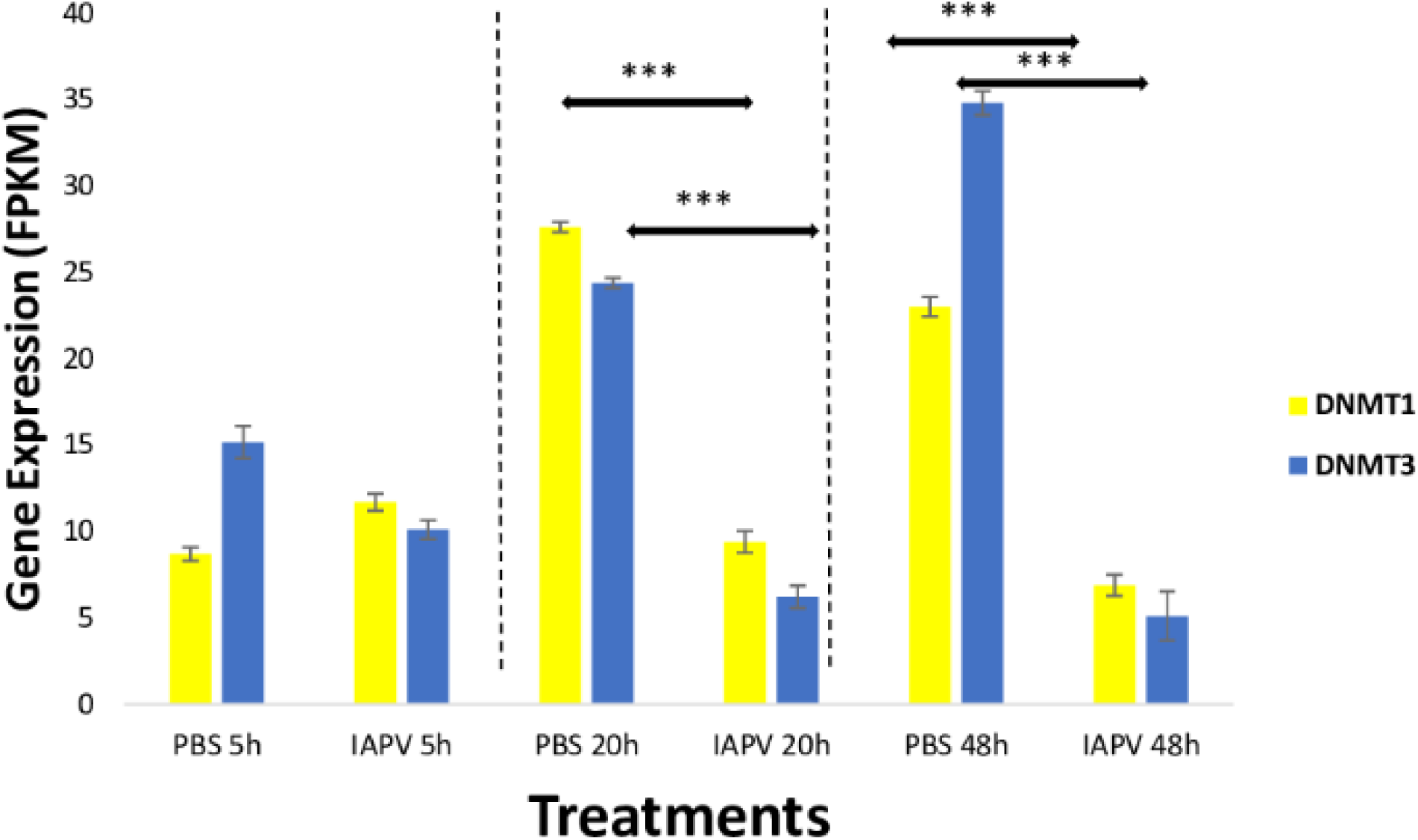
Gene expression patterns (mean ± SE) of the major DNA Methyl-Transferases (DNMTs). Both DNMT1 (GB47348) and DNMT3 (GB55485) showed significant down-regulation when comparing sham control (N = 3) and IAPV samples (N=3) after 20 h and 48 h after IAPV infection respectively. ***: p < 0.001. False Discovery Rate (FDR) < 0.05.

### Changes of DNA methylation after viral infections and comparative methylomic analysis

Total methylated 5mC sites in CpG, CHG, and CHH settings were 1,955,471; 878,838; and 1,433,905 respectively (Figure 7). The ratio of methylated CpG sites versus total CpG sites was highest in 5h PBS control (10%), followed by 5h IAPV post-infection samples, 20h PBS control, 20h IAPV post-infection samples, 48h control, and lowest in 48h IAPV post-infection samples (about 4%) (Figure 6). The whole genome-wide Pearson correlation matrix for CpG base profiles across all samples at each time point (5h, 20h, and 48hr post-infection) are listed in supplemental figure 8.

**Fig. 7:**
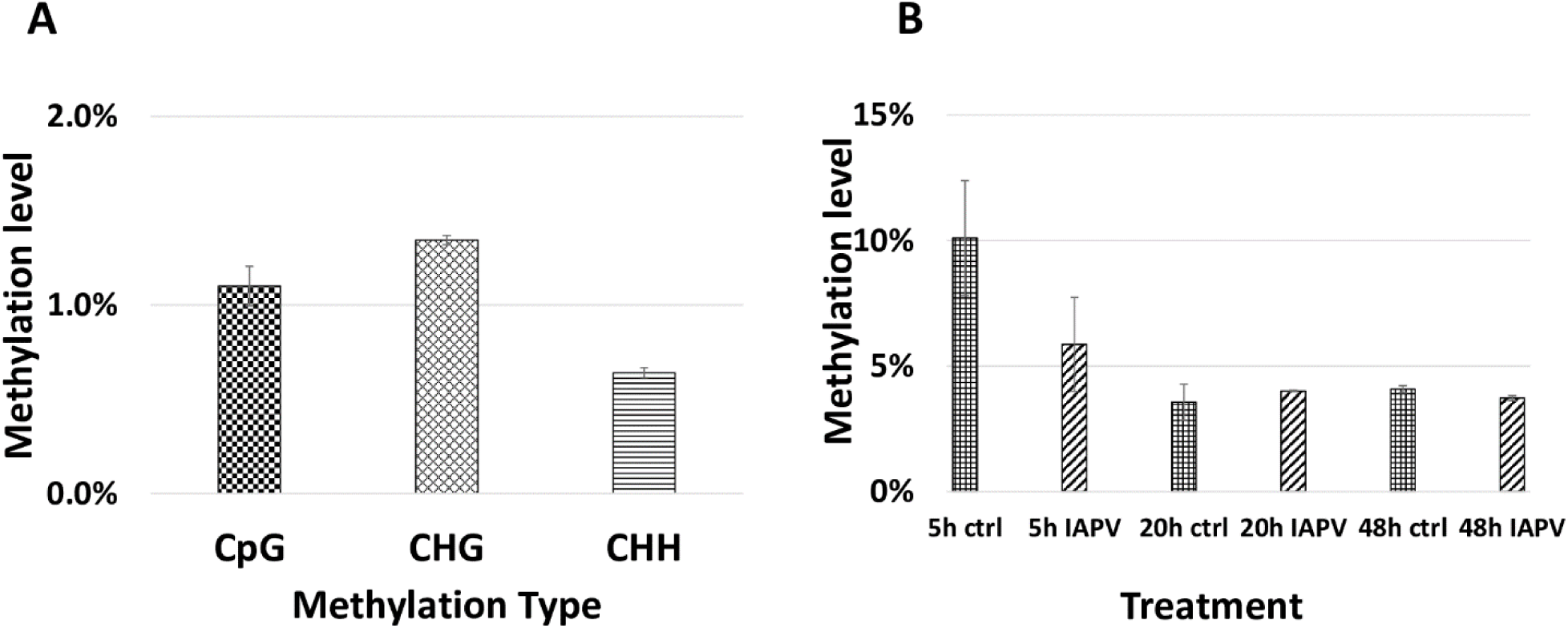
Genome wide methylation patterns. A. Total methylated 5mC sites in each group (CpG, CHG, and CHH) are shown. B. Methylated CpG sites/Total CpG sites are shown for each treatment and time point.

We identified 523, 5, and 18 differentially methylated regions (DMRs) between IAPV and control groups in the 5h, 20h, and 48h post-infection samples, respectively (Supplemental Table 3 for a detailed gene list). Significant overlap of DMRs existed between 5h and 20h (p<0.05) and 5h and 48h (p<0.0001) comparisons (Supplemental Table 4).

Our comparison of our DMR lists with the previously published immune genes of Evans et al. [41] revealed 5 and 1 gene were overlapping with the DMRs at 5h and 20h/48h post-infection, respectively (Figure 4B, Supplemental Table 3 & 4). Comparison of our data with IAPV-induced methylation changes in adult honey bees of Galbraith et al [24] revealed 14 and 1 genes were overlapped with DMRs at 5h and 20h/48 h (Figure 4D, for specific gene names see Supplemental Table 3 & 4).

We examined the genome-wide Pearson correlation matrix for CpG base profiles across all samples at each time point. Overall samples after 5h infection have the highest correlation (0.72-0.87), and samples after 48hr infection have the lowest correlation (0.69-0.73); specific coefficients are listed in Figure 8).

### Differential DNA methylation leads to differential gene expression later in several immune pathways

To look into the potential association between gene expression and CpG methylation, we studied whether genes that showed early changes of CpG methylation also exhibited significant changes of gene expression later. A list of 38 immune genes that were hypo- or hyper-methylated at the first time point (5h post-IAPV infection), were found differentially expressed later (48h post IAPV infection) (Supplemental Table 8). These cellular pathways included the MAPK (mitogen-activated protein kinase) pathway, JAK/STAT (Janus kinase/signal transducer and activator of transcription) pathway, Hippo signaling pathway, mTOR (mammalian target of rapamycin) signaling pathway, TGF-beta (transforming growth factor-beta) signaling pathway, ubiquitin mediated proteolysis, and spliceosome. Interestingly, we detected 12 genes from this group with differentially spliced patterns with a significant statistical q value (Table 2).

**Table 2.**
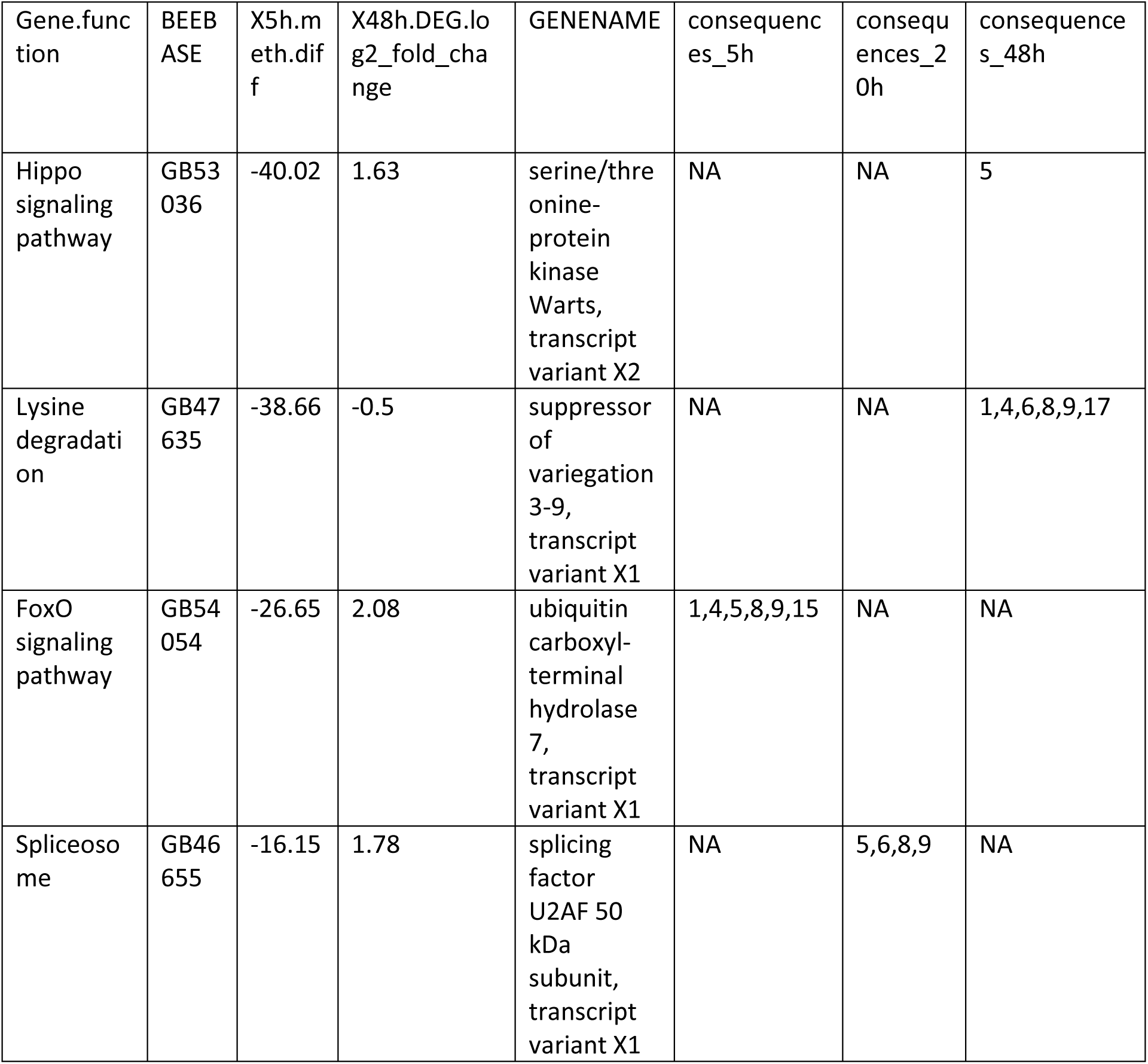
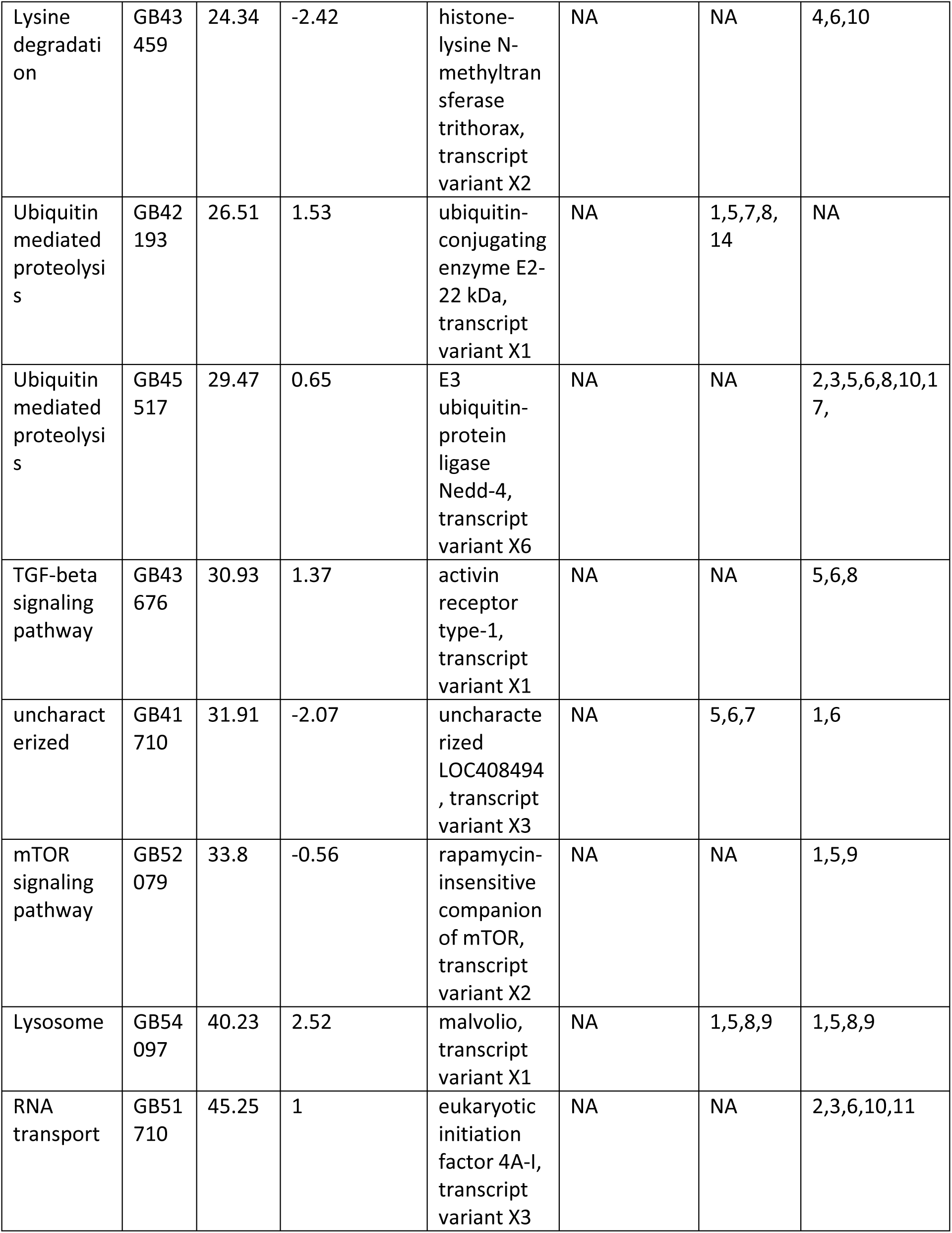
A list of twelve immune genes that were hypo- or hyper-methylated first post 5h of viral infection, then gene expression and splicing patterns changed significantly post 48h of viral infection (q < 0.05). Gene.function (column 1) showed immune pathways involved. BEEBASE (column 2) showed the gene ID in BEEBASE. X5h.meth.diff (column 3) showed changes of DNA methylation level of the gene/region when comparing post 5h IAPV infection samples and control samples. X48h.DEG.log2_fold_change (column 4) showed fold change of gene expression in log2 scale when comparing post 48h IAPV infection samples and control samples. GENENAME (column 5) showed gene names. Consequences_5h, consequences_20h & consequences_48h (column 6-8) are the detected categories of alternative splicing listed in the first column of Table 1. Details of gene IDs and q values are in Supplemental Table 8.

| Gene.function | BEEBASE | X5h.meth.diff | X48h.DEG.log2_fold_change | GENENAME | consequences_5h | consequences_20h | consequences_48h |
| --- | --- | --- | --- | --- | --- | --- | --- |
| Hippo signaling pathway | GB53036 | -40.02 | 1.63 | serine/threonine-protein kinase Warts, transcript variant X2 | NA | NA | 5 |
| Lysine degradation | GB47635 | -38.66 | -0.5 | suppressor of variegation 3-9, transcript variant X1 | NA | NA | 1,4,6,8,9,17 |
| FoxO signaling pathway | GB54054 | -26.65 | 2.08 | ubiquitin carboxyl-terminal hydrolase 7, transcript variant X1 | 1,4,5,8,9,15 | NA | NA |
| Spliceosome | GB46655 | -16.15 | 1.78 | splicing factor U2AF 50 kDa subunit, transcript variant X1 | NA | 5,6,8,9 | NA |
| Running title: Methylated genes involved in honeybee IAPV-infection |  |  |  |  |  |  |  |
| Lysine degradation | GB43459 | 24.34 | -2.42 | histone-lysine N-methyltransferase trithorax, transcript variant X2 | NA | NA | 4,6,10 |
| Ubiquitin mediated proteolysis | GB42193 | 26.51 | 1.53 | ubiquitin-conjugating enzyme E2-22 kDa, transcript variant X1 | NA | 1,5,7,8,14 | NA |
| Ubiquitin mediated proteolysis | GB45517 | 29.47 | 0.65 | E3 ubiquitin-protein ligase Nedd-4, transcript variant X6 | NA | NA | 2,3,5,6,8,10,17, |
| TGF-beta signaling pathway | GB43676 | 30.93 | 1.37 | activin receptor type-1, transcript variant X1 | NA | NA | 5,6,8 |
| uncharacterized | GB41710 | 31.91 | -2.07 | uncharacterized LOC408494, transcript variant X3 | NA | 5,6,7 | 1,6 |
| mTOR signaling pathway | GB52079 | 33.8 | -0.56 | rapamycin-insensitive companion of mTOR, transcript variant X2 | NA | NA | 1,5,9 |
| Lysosome | GB54097 | 40.23 | 2.52 | malvolio, transcript variant X1 | NA | 1,5,8,9 | 1,5,8,9 |
| RNA transport | GB51710 | 45.25 | 1 | eukaryotic initiation factor 4A-I, transcript variant X3 | NA | NA | 2,3,6,10,11 |

### Comparative methylomic analyses of immune genes with other pathogen infection

The comparison with a previously published list of 186 immune genes from Evans et al (2005) [41], showed that 5 (*GB45248, GB46478, GB55483, GB51399*, and *GB41850*) and 1 (*GB46478*) genes overlapped in our 5h and 48h DMR lists, respectively (Supplemental Table 3). There was no significant directional overlap of the list of immune genes and our hyper- or hypo-regulated DMRs at 5h, 20h, and 48 h (Supplemental Tables 3 & 4). Comparison of our data with IAPV-induced gene expression changes in adult honey bees in Galbraith et al [24] revealed 19, 1 (*GB51998*), and 1 genes (*GB42022*) overlapped, indicating no significant directional overlap of our hyper- or hypo-regulated DMRs at 5h, 20h, and 48 h (Supplemental Tables 3 & 4). When compared to immune genes associated with anti-viral responses in *Drosophila* [52], 20, 1 (*GB17743*), and 1 genes (*GB15242*) overlapped in our 5h, 20h, and 48h DMR lists (Supplemental Table 5).

### Motif

To reveal the potential regulation of CpG methylation, we examined the 6-mer motif of each CpG site, including the 2 nucleotides upstream and downstream of each CpG site. Analysis of 6-mer motifs from 5h DMRs showed no common pattern of the motif (Supplemental Figure 11). Similarly, analysis of motifs from 20h DMRs containing CpG sites revealed little signal beyond the selected central CpG sites in the middle of 6-mer motifs. However, DMRs from 48h post-IAPV infection tend to have a T or C following the central CpG (Supplemental Figure 11).

## Discussion

To test our hypothesis that viral infections not only affects gene expression profiles but also DNA methylomic marks in honey bee pupae, we compared the complete transcriptome and methylome responses across three distinct phases (5h, 20h, and 48h) of a lethal IAPV infection. While methylation is most affected early during the infection, differential gene expression increases over time and severity of the infection. Many up- and down-regulated genes are involved in RNA processing and translations, proteolysis, and metabolic processes (Figure 3). A collection of genes have differential transcript usage or alternative splicing post IAPV infections (Figure 5). When compared to differentially expressed genes from adult honey bee and fruit fly post viral infection, we have discovered significant overlap of our IAPV-responsive genes with immune genes from these other studies [41].

The pupal stage is a critical phase for metamorphosis and an important temporal window for *Varroa* mite parasitism. Our study, which incorporates and confirms the separate observations of adult or larval honey bees [24, 44, 47, 53], includes new observations that provide a more complete understanding of how the pupae respond to lethal viral attack. Previous studies showed that honey bee pupae can respond to disease with increased immune gene expression to decrease the success of parasites and pathogens [42, 54]. Our results reveal that rapid epigenetic changes in response to viral infection precede and presumably trigger profound and widespread transcriptome responses in honey bees. A combination of both transcriptomics and methylomics at multiple time points can reveal the role of and the relation between 5mC methylation and gene expression profiles under viral infection of insects [55].

Alternative splicing in response to viral infection can affect subsequent gene expression through several mechanisms [56]. Our results of differential transcript usage or alternative splicing indicated the viral infection can cause large changes in alternative splicing of the bee hosts, which brings a new perspective on molecular mechanisms of IAPV virus pathogenesis. It is known that RNA virus can alter host mRNA splicing [57, 58]. Previous research revealed that DNA methylation is linked to regulate alternative splicing in honeybees via RNA interference knocking down DNMT3 [5].

The epigenetic changes we report here are interesting because the 5mC methylome profile indicated a large amount of DMRs after 5h IAPV infection. However, DMR number dramatically dropped after 20h and 48h of infection. This phenomenon may be explained by the cell death and apoptosis after severe infections after 20h and 48h. We propose a possible explanation is that 5mC changes as a primary host-defense mechanism responding to viral infection at the onset. During the early stage of viral infection, 5mC methylation may be the first to react leading to molecular changes genome-wide, potentially interacting with the transcription process such as transcription factor binding. The mechanism on how to induce abnormal methylation at the molecular and cellular levels by viral infection is yet to be determined. A few potential explanations are that pathogenic infection may lead to cellular proliferation and inaccuracy in epigenetic processes, or molecular signaling pathways involved in the infection response affect these epigenetic processes. Previous studies showed that viral infection can change 5mC methylation in human cancer [59, 60]. Other studies also showed that changes in epigenetics marks can affect immune responses, cell-cycle checkpoints, cell death, and cell fate [59, 61].

In honey bees, some evidence links 5mC methylation marks to gene regulation [5, 12, 23, 62, 63] but limited reports showed 5mC methylation are associated with change of gene expressions [21]. The data here indicates that 38 of the initially differentially methylated genes subsequently exhibit significant changes of gene expression (Table 1), and 12 of these genes displayed significant changes in their splicing events. In general, the epigenetic control of 5mC methylation and gene expression has been reported for plants and mammals [64, 65], but the function of 5mC methylation in the regulation of genes in insects is still not clear [5, 62]. One hypothesis is that DNA methylation is involved in cell signaling or involved in the regulation of co-transcription via transcription factors [66-68].

DNA methyl-transferases 1 and 3 are critical enzymes for a functional methylation system. Our data show the IAPV infection reduced the gene expression of these two key enzymes over time. This finding is consistent with previous reports showing that viral infection can change the expression of DNMTs in animal and cell studies [69, 70]. The significant fold change of these enzymes is moderate but sufficient to cause dramatic changes in the global DNA methylome, as indicated in previous studies [5, 11, 12]. The DNA methylation changes in turn can have further consequences for widespread gene expression patterns. We found more DMR post 5h IAPV infection, when the DNMTs are not yet differently expressed. Possible explanation is that the initial viral infection induced the methylation changes at the initial stage by other potential molecular mechanism such as histone modifications [71, 72]. DNMTs may be regulated to restore homeostasis which may explain why fewer DMRs at the later stages [73].

At every time-point, we identified numerous differentially expressed genes that confirmed the profound consequences of IAPV infection [51]. Such effects can certainly be caused by cell expansion or loss, rather than direct effects on gene activity. However, the systemic consequences for the health and function of the organism may be similar: the level of immunity might drop due to declining cell numbers or lowered gene activity per cell. Molecular pathways from differentially expressed and methylated genes indicated that the MAPK signaling pathway, Jak/STAT signaling pathway, Hippo signaling pathway, mTOR signaling pathway, TGF-beta signaling pathway, ubiquitin mediated proteolysis, and spliceosome are critical to respond to viral infections. Both MAPK and Jak/STAT signaling pathway are immune pathways with antiviral immune response in honey bees and dipterans [24, 28, 40, 43, 47, 74, 75]. The Hippo signaling pathway is highly conserved from insects to mammals, which regulates cell death and differentiation [76]. Previous study also showed differentially expressed genes regulated by IAPV in honey bee brood are involved in the mTOR, TGF-beta pathways, and ubiquitin mediated proteolysis [47].

In addition, our data showed that IAPV infection can induce *Defensin-1*, a key antimicrobial peptide responding to viral infections [40, 54]. P450 genes are known as responding to oxidative stress in *Drosophila* during larval development [77]. In honey bees, P450s are involved in xenobiotic detoxification [78, 79]. From a wider perspective, Chikungunya virus infection specifically activates the MAPK signaling pathways in humans [80] and is a stress-responsive part of the intestinal innate immunity of *C. elegans* [81]. Recent report showed that MAPK may play a role in regulating certain P450 genes [82]. On the other hand, IAPV infection presumably causes major deregulation of the cellular homeostasis through the cooption of the cellular machinery to replicate itself [51]. Thus, deregulation of ribosome biosynthesis (Fig. 3) and major energetic pathways is not surprising. Interestingly, transcriptomic analyses of Colony Collapse Disorder, which had been associated with IAPV [44], have shown similar immune responses [79, 83].

The methylomic profile of pupae 48h after IAPV infection indicated genes involved in anti-parasitoid immune response [84, 85] to be affected. One of these genes that overlaps with a previous genomic study of honey bee immune genes [41] is Peroxin 23, a protein involved in peroxisomal protein import and associated with autophagosome formation in *Drosophila* muscles and central nervous system [86, 87]. Compared to a previous report of the 5mC methylomic profiles of honey bee adults infected with IAPV [24], our analysis revealed only one overlapping gene (GB51998), which is related to ATP binding, circadian regulation of gene expression systems [88], and muscle morphogenesis and function in *Drosophila* [89].

The list of identified DMRs linked to genes associated with ATP binding, phagosome, fatty acid degradation, RNA transport, and RNA degradation. Lipid metabolism, signaling, and biosynthesis are related to virus-host interactions [90]. Fatty acid degradation and metabolism also responded to cold stress in other insects and might indicate increased energetic demands of stressed organisms [91]. Alternatively, IAPV may remodel the host cells by hijacking the lipid signaling pathways and synthesis process for its own replication, similar to virus manipulation of host RNA replication and transport mechanisms [49, 51].

Our study only represents a first step in the understanding of the intricate interplay of viruses and their honey bee host’s physiology. Future research needs to characterize the temporal transcriptome trends in more detail and tissue-specificity. Pupae are particularly relevant but also challenging because the ongoing metamorphosis entails drastic hormonal and presumably transcriptional changes even in the absence of disease. The discovered functional pathways need to be functionally investigated, particularly the putative 5mC methylation control of immune genes.

## Materials and Methods

### Bee samples and Virus preparation

All honey bee samples were acquired from unselected experimental hives in the research apiary of the University of North Carolina Greensboro. A previously prepared and characterized IAPV solution in PBS was used for inoculations[51]. White-eyed pupae were either injected with PBS solution as control treatment or with 10^4^ genome copies of IAPV in 1.0 μl of the viral suspension per pupa. Pupae were maintained on folder filter papers in a sterile lab incubator at 32 °C and 60%R.H. After 5h, 20h, and 48h of injections, each individual pupal sample was flash-frozen in liquid nitrogen and then stored in a -80 °C freezer. Figure 1 illustrates the experimental design.

### RNA-Seq library preparation and sequencing

Treatment control (PBS-injection) and IAPV-infected pupae were compared across three time points (5h, 20h, and 48h). Three pupae were collected for each treatment and each time point for a total of 18 samples. Dual extraction of nucleic acid (RNA and DNA) from each pupa was carried out using the MasterPure dual extraction kit (Epicentre, MC85200). Briefly, whole pupae were individually homogenized and lysed first by using a micropestle. The lysed cells were mixed with extraction buffer and centrifuged to remove proteins and undesired debris of the cell. Subsequent RNA and DNA fractions were treated with DNase and RNase accordingly. RNA-seq library preparation and sequencing were performed by the Genomic Sciences Laboratory of the North Carolina State University. Total RNA from each pupa was used to generate tagged Illumina libraries, using the NEBNext^®^ RNA Library Prep Kit (New England Biolab). All RNA-seq libraries were sequenced in two flow cells on the Illumina Next-Seq 500 (paired-end 150bp length reads).

### BS-Seq library preparation and sequencing

Bisulfite (BS) conversion of the genomic DNA from the above-mentioned dual extraction of each sample was carried out by using the EZ DNA Methylation-Lightning Kit (Zymo Research, D5030, Irvine, CA). BS-converted DNA (100 ng) of each sample was used for preparing the BS-seq library via the TruSeq DNA Methylation Kit (Illumina, EGMK91324, San Diego, CA). The quality of the libraries was checked using Qubit assays (Q32850, Life Technologies) and with the 2100 Bioanalyzer (Agilent Technologies, Inc.). The pooled 18 libraries were run on two replicate lanes of the Illumina NextSeq 500 (150 bp paired-end reads) by the Genomic Sciences Laboratory at NCSU.

### Bioinformatic analyses

Overall quality control analysis of all resulting data was carried out by using FastQC (http://www.bioinformatics.babraham.ac.uk/projects/fastqc/). Raw data were trimmed by Trimmomatic (java -jar PE -phred33 ILLUMINACLIP:TruSeq3-PE.fa:2:30:10 MINLEN:36). TopHat2 v.2.0.12 [92] was used to align trimmed RNA-seq reads to the *Apis mellifera* 4.5 reference genome. General alignment parameters were chosen to allow three mismatches per segment, 12 mismatches per read, and gaps of up to 3bp (tophat --read-mismatches 12 --segment-mismatches 3 --read-gap-length 3 --read-edit-dist 15 --b2-sensitive) to minimize the possibility that small sequence differences of our samples with the reference genome would bias expression estimates. These settings resulted in the successful alignment of 35% of raw reads (117–143 million per sample). Reconstructed transcripts were aligned with Cufflinks (v2.2.1) to the *Apis mellifera* OGS 3.2 allowing gaps of up to 1 bp (cufflinks --overlap-radius 1 --library-type fr-firststrand). To annotate promoters and transcription starts, we employed the ‘cuffcompare’ command using OGS3.2 gff file (cuffcompare -s -CG -r).

Cuffdiff2 (v.2.1.1) [92] was used to test for differential expression between IAPV and control groups (n=3 pupae treated as biological replicates at each of the 3 different time points) with the multiple mapping correction and a false discovery rate threshold set at 0.05. All three IAPV groups (IAPV_5h, IAPV_20h, and IAPV_48h) were compared to their corresponding sham-control groups (PBS_5h, PBS_20h, PBS_48h) to identify differentially expressed genes (DEGs). The data were explored and visualized with CummeRbundv.2.0.0 (http://compbio.mit.edu/cummeRbund/) and custom R scripts (Rsoftwarev.2.15.0, http://www.r-project.org/). Differentially expressed genes with similar expression temporal dynamics were analyzed by cufflinks in R. Total of output of gene numbers is set to be 10 (for example, consider gene GB49890 (cyp6as5), we can find nine other genes in the database with similar expression patterns, then plot the expression patterns in Supplemental Figure 7).

Whole genome BS-Seq analysis was carried out by trimming the sequences using trim_galore (version 0.4.1) (http://www.bioinformatics.babraham.ac.uk/projects/trim_galore/) (trim_galore ../file1_R1_001.fastq.gz ../file1_R2_001.fastq.gz -q 20 --paired --phred33 -o). The Bismark tool [93] (bismark --bowtie2 --path_to_bowtie -p 8 path_to_genome_files/ --1 /file1 --2 /file2) was employed for whole genome alignment. Brief procedures included Genome Conversion, Genome Alignment, Deduplication, and Methylation calls. Differentially methylated regions and genes (DMRs) were produced by methylKit in R (version 1.3.8) with percent methylation difference larger than 10% and q < 0.01 for the difference [94].

The reference genome and official gene set were the same as described before. The sequencing raw data are published at the NCBI database (Sequence Read Archive (SRA) submission: SUB3404557, BioProject: PRJNA429508). All codes for the bioinformatic analyses are online (https://www.researchgate.net/publication/343237015_RNAseq_and_BSseq_codes_for_public).

To determine overlap between DEG lists, either up- or down-regulated genes were compared between different time points. This directional overlap analysis avoids artifacts that can confound undirected DEG comparisons [95]. The analysis was compared focusing on DEGs with a more than 8-fold difference, and with DMRs (either hyper- or hypo-methylated). All overlap analyses were performed with hypergeometric tests to calculate the p-value for overlap based on the cumulative distribution function (CDF) of the hypergeometric distribution as published previously [5]. GO Mann-Whitney U tests were carried out on the gene expression data using codes published in Github (https://github.com/z0on/GO_MWU). These analyses search for patterns of up- or down-regulation in genes associated with particular GO-terms based on their signed, uncorrected p-values regardless of significance thresholds [96].

For differentially transcript usage or spliced genes, transcript-level counts were quantified using Salmon1 v1.2.1[97] against the A. mellifera HAv3.1 annotation. Viral RNAs were included in the transcriptome used for alignment but were omitted from isoform analysis. Salmon results were imported directly into the IsoformSwitchAnalyzeR R package[98, 99], which normalizes transcript counts based on abundances and uses the TMM method to adjust effective library size. The version of IsoformSwitchAnalyzeR used was 1.11.3, obtained from GitHub commit 5a7ab4a due to a bug in version 1.11.2 on Bioconductor. Although CDS were annotated in the GFF file for A. mellifera, these caused problems in IsoformSwitchAnalyzeR due to inclusion of stop codons, as well as some transcripts having no annotated CDS or CDS that were obviously incorrect. Therefore, the analyzeORF function was used to predict open reading frames *de novo* from the transcript sequences, which in the vast majority of cases resulted in agreement with the published annotation. Nucleotide and amino acid sequences for all transcripts for significant genes were then exported for analysis with external tools: CPAT9 was used to determine whether a transcript was likely to be coding [100]. It was run from the webserver using the *Drosophila* (dm3, BDGP release 5) assembly. A coding probability of 0.7 was used as the cutoff for classifying a transcript as coding or non-coding. Pfam10 [101] version 32.0 was used to predict protein domains, using pfam_scan v1.6 run on the Biocluster.

### Statistical analysis of differential transcript usage

A design matrix was constructed in which the condition was the combination of treatment and time point (six conditions total), and DWV infection (determined by alignment of reads to the DWV mRNA) was included as a covariate. Sample degradation was excluded as a covariate as it introduced singularity into the model at 48h. The contrasts tested were IAPV_5h - PBS_5h, IAPV_20h - PBS_20h, and IAPV_48h - PBS_48h. The preFilter function in IsoformSwitchAnalyzeR was used with default settings to filter transcripts and genes. Genes were removed if they only had one isoform or if they did not have at least 1 RPKM in at least one condition. Transcripts were removed if they had no expression in any condition. After filtering, 15,808 isoforms belonging to 4,667 genes remained.

Differential transcript usage (DTU) was assessed using DEXSeq from within IsoformSwitchAnalyzeR [99, 102, 103]. Genes were only retained for further analysis if they had an FDR < 0.05 for DTU, and if isoform usage (proportion of counts from a gene belonging to a given isoform) changed by at least 0.1 for at least two isoforms in opposite directions (i.e. the isoforms “switched”).

### Enrichment of overlapping genes among different studies

To test whether our DEGs and DMRs had significant overlap with known immune genes, these lists were compared to published immune genes [41] using hypergeometric tests. Additionally, a directional overlap analysis was performed on our DEGs (separated by up- or down-regulation) with a previous RNA-seq study of IAPV infection in adult honey bee workers [24].

To identify potential motifs that are targeted for epigenetic modification after IAPV infection, differentially methylated regions (DMR) were analyzed post 5h, 20h, and 48h IAPV infection generating 6-mer sequence logos with the online tool weblogo (https://weblogo.berkeley.edu).

## Acknowledgements

The National Research Council Postdoctoral Fellowship, USDA NIFA Evans Allen fund (NI191445XXXXG002), NSF HBCU-UP grant (1900793), and Illumina Go Mini Challenge Grant supported H.L.-B. Major research funding was from the U.S. Army Research Laboratory grant W911NF-04-D-0003 to OR, DRT, and MKS. We thank Karl M. Glastad (University of Pennsylvania), Marce Lorenzen (NCSU), Andrew Petersen (NCSU), and HPCBio at University of Illinois on bioinformatics consulting, Karmi Oxman and Susan Balfe for the graphic designs, and Conrad Zagory, Jr. at CSU for critical reviews on the English writing.

## Figure legends

Table 1. A list of twelve immune genes that were hypo- or hyper-methylated first post 5h of viral infection, then gene expression and splicing patterns changed significantly post 48h of viral infection (q < 0.05). Tss (Transcription start site), Tts (Transcription termination site), ORF (Open Reading Frame), UTR (Untranslated Region), NMD (Nonsense-mediated decay), IDR (intrinsically disordered regions).

Table 2. A list of twelve immune genes that were hypo- or hyper-methylated first post 5h of viral infection, then gene expression and splicing patterns changed significantly post 48h of viral infection (q < 0.05). Details of gene IDs and q values are in Supplemental Table 8.

